# Invasion and secondary site colonization as a function of in vitro primary tumor matrix stiffness

**DOI:** 10.1101/2022.07.22.500985

**Authors:** Lekha Shah, Ayşe Latif, Kaye J. Williams, Elena Mancuso, Annalisa Tirella

## Abstract

Increased breast tissue stiffness is correlated with breast cancer risk and invasive cancer progression. However, its role in promoting bone metastasis, which shares a large burden of breast cancer deaths, has not yet been understood. To better understand the cause-effect relationship of tissue stiffness on breast cancer’s metastatic potential, we fabricated three-dimensional (3D) models to mimic breast and bone tissue *in vitro*. Based on our previous work, we used alginate-based hydrogels allowing precise control over stiffness and composition of extracellular breast tissue matrix; and 3D printed poly-caprolactone (PCL)-composite scaffolds to mimic the bone. The latter were further modified by promoting bone-ECM deposition using Saos-2 cells. After a decellularization step, PCL scaffolds were assembled with alginate-gelatin hydrogels and a novel breast-to-bone *in vitro* model was established. It was observed that increased stiffness of hydrogel resulted in higher migration and invasion capacity of MDA-MB 231 cells. Additionally, PTHrP and IL-6 expression, both of which are implicated in bone metastasis, were higher when cells from stiff hydrogels were cultured in bone/PCL scaffolds. These breast-to-bone in vitro models pose as a novel non-animal technology to pave the way for incorporating important tissue microenvironmental factors of the disease physiology (e.g. tissue stiffness) and emerge as promising future platforms for monitoring metastatic disease phenotypes and therapeutic efficacy.

## 1. Introduction

Dynamic communication between cells and extracellular matrix (ECM) governs tissue structure and function. However, this homeostasis is dysregulated in cancer wherein tumor cells and cancer associated fibroblasts actively remodel the ECM, leading to increased deposition, crosslinking and linearization of fibrillar components such as collagen ^1,2^. This leads to a change in biomechanical and biochemical nature of the ECM with modifications of matrix stiffness and tissue architecture, as well as cell adhesion motifs ^3^. Tumor breast tissue stiffness is reported to be 3-6 times stiffer than normal tissue, and is used clinically in breast cancer diagnosis ^4^. Due to their long-term nature which are shared across patients, tissue stiffening and ECM changes are used as clinical diagnostic technique than other markers, having a significant importance in solid cancer progression ^5^. As a matter of fact, ECM modifications are proved to be implicated in cell fate ^6^, cancer stemness ^7^, cancer progression^3^, metastasis ^8^ and therapeutic response in tumor tissue ^9,10^.

Studies have investigated effects of stiffness on breast cancer progression in vitro. For example, a stiffness increase from 0.2 kPa to 5 kPa in peptide crosslinked polyacrylamide gels was observed to perturb mammary acini formation by inducing unhindered growth and polarity loss in mammary epithelial cells ^11^. Stiff matrices (i.e. > 3 kPa) were also observed to affect genome wide expression changes in non-malignant breast epithelial cells and found to develop malignant phenotype by altering chromatin accessibility of genes ^12^. However, very few studies illustrate effects of primary tumor stiffness, and ECM properties, on invasion and metastasis to a secondary site. Many in vitro studies model metastatic sites only ^13,14^, and none has been reported to explore the connection between primary and the secondary site in the same model.

Bone is the most common site for breast cancer metastasis (∼60-70%) followed by lung, liver and brain ^15^. Breast cancer metastasis to a secondary site like bone involves a change in ECM composition, which in contrast to breast’s collagenous matrix, consists of ∼60% inorganic matrix (majorly hydroxyapatite/HA) and ∼40% organic matrix (majorly collagenous) ^16^. Highly vascular trabecular bone, which is a preferred site for bone metastasis ^17^, has high porosity (average ∼ 40%) and a heterogenous tissue stiffness that ranges from 4-80 MPa ^18^, whereas normal breast and tumor tissue are comparatively dense with tissue stiffness ranging from 3-16 kPa ^4^. When a subpopulation of circulatory breast tumor cells encounters bone ECM, they start a cascade of bone metastatic cycle. An important first step of this cycle is when tumor cells release Parathyroid Hormone related Protein (PTHrP) in response to bone’s increased extracellular calcium ^19^ and Transforming Growth Factor-β (TGF-β) ^20^. Tumor secreted cytokines such as Interleukin-6 (IL-6) ^21^ along with PTHrP ^22^ instructs osteoblasts to release Receptor Activator of Nuclear factor Kappa-B? Ligand (RANKL) which consequently stimulates osteoclast activity. This forms osteolytic bone lesions that further releases trapped calcium and TGF-β to induce cancer cell signaling, hence promoting the cycle.

Increase of primary matrix stiffness is recorded in many solid tumors, and cells mechanically conditioned to such stiffness maintain their phenotype even after removal of mechanical stimuli ^23,24^. This is of particular relevance at later stages of tumor progression i.e., invasion and metastasis, which involves leaving the primary tumor site and invading spaces with different ECM properties. Until now, no answer on how primary tumor ECM properties can condition cells and dictate their response to a secondary site ECM has been provided. In this scenario, 3D in vitro models can offer an opportunity to model ECM changes with required precision, as well as decouple biomechanical and biochemical components to understand their individual contribution. We previously characterized alginate-gelatin hydrogels to mimic normal breast tissue stiffness (2 kPa) and tumor tissue stiffness (6-10 kPa), varying the concentration of gelatin to change composition and density of hydrogels ^25^. We have also used composite poly-ε-caprolactone (PCL) scaffolds to mimic the bone tissue (i.e. PCL scaffolds containing inorganic compounds such as hydroxyapatite (HA), strontium HA (SrHA) and barium titanate (BaTiO_3_) with stiffness and porosity matching the values of trabecular bone ^26,27^, being 40-55 MPa and 35-45% respectively.

In this study, key aspects of breast and bone tissue ECM were engineered, namely matrix composition, stiffness, density, porosity, and architecture to study breast-to-bone metastasis. In particular, to increase biocompatibility and physiological relevance of composite PCL scaffolds, Saos-2 were cultured within these scaffolds, and decellularization procedures were optimized to retain mineralized deposited ECM (named as *biohybrid PCL scaffolds*) and used as bone secondary metastatic site.

Effect of primary tumor matrix stiffness on human breast cancer cells (MDA-MB 231) was analyzed testing various aspects of the metastatic cascade like cell adhesion, migration, 3D invasion and secondary site metastasis through in vitro assays. It was observed that there is a proportional increase of migration, invasion and osteolytic factor expression (PTHrP and IL-6) in MDA-MB 231 cells with an increase in stiffness of hydrogels used to pre-condition these cells.

This study aimed at modelling in vitro important properties of the tumor microenvironment and allowing the detection of cellular events directing biological processes (e.g. extravasation, migration, invasion) and metastatic onsets. In particular, the study focused on: 1) effect of breast tissue stiffness on invasive potential and 2) mechanism of bone invasion. We showed that high primary tumour stiffness can induce high osteolytic factor (i.e. PTHrP) production in breast cancer cells once they reach the in vitro bone tumour microenvironment. The proposed models are designed to deepening understanding of the complex metastatic cascade, with potential impact on the identification of new therapeutic modalities (e.g. mechano-therapeutics) and accelerate their translation to the clinic.

## 2. Methodology

### 2.1 General Cell culture

Human breast adenocarcinoma cell line MDA-MB 231 (HTB-26, ATCC) and breast cancer bone homing cell line MDA-IV (kindly provided by Prof I Holen, University of Sheffield, Sheffield, UK) were cultured in Dulbecco’s Modified Eagle Medium (DMEM, D6546, Sigma-Aldrich, UK) supplemented with 1% (v/v) L-glutamine (G7513, Sigma-Aldrich, UK), 10% (v/v) Fetal Bovine Serum (FBS, F9665, Sigma-Aldrich, UK) and 1% (v/v) penicillin-streptomycin (P4333, Sigma-Aldrich, UK). Human osteosarcoma cell line Saos-2 (kindly provided by Dr O Tsikou, The University of Manchester, Manchester, UK) were cultured in McCoy’s 5A media (M9309, Sigma-Aldrich, UK) supplemented with 15% (v/v) FBS and 1% (v/v) Penicillin-Streptomycin.

All cell lines were tested negative for mycoplasma contamination by Mycoalert mycoplasma detection kit (LT07-318, Lonza) prior use. Unless otherwise specified, all cell culture experiments were performed in a humidified 5% (v/v) CO_2_ air atmosphere at 37 °C in complete medium. Cells were used from passage 9 to 25.

### 2.2 3D cell culture: encapsulation in alginate-gelatin hydrogel beads

#### 2.2.1 Preparation of hydrogel precursors and crosslinking solutions

Four different combinations of alginate and gelatin hydrogel precursor solutions (So-L, So-H, St-L and St-H) were selected and prepared as described in our previous study ^25^ to match stiffness and composition of normal and tumor breast cancer tissue (Table 1). Briefly, sodium alginic acid (G/M ratio of 0.7, Pro-Alg, Chile) was dissolved in HEPES buffered saline (HBS) at a concentration of 3% and 6% (w/v). Obtained alginate solutions (aq.) were sterile filtered with 0.22 µm PES filter. Gelatin type A (G1890, Sigma-Aldrich, UK) was dissolved in HBS, at a concentration of 2% and 6%. Gelatin solutions (aq.) were sterile filtered with 0.45 µm PVDF filter. Hydrogel precursor solutions were prepared by mixing (5 min, RT) different combination of alginate and gelatin solutions at selected concentrations with a 1:1 volume ratio (final hydrogel composition is reported in Table 1). Calcium chloride (CaCl_2_, C/1400/53, Fischer scientific, UK) solutions were prepared in deionized water at concentrations of 100 mM, 200 mM and 300 mM. Each solution was sterile filtered with 0.22 µm PES filter prior use and stored at 4 °C.

**Table 1:** Hydrogel composition of the four selected alginate-gelatin hydrogels varying compressive moduli (stiffness) and concentration of gelatin (adhesion ligand content, hydrogel density). Sample ID are named with a combination of their properties: Soft (So, stiffness < 3 kPa), Stiff (St, stiffness > 6 kPa), low adhesion ligand (L, 1% w/v gelatin), high adhesion ligand (H, 3% w/v/ gelatin).

| Sample ID | Hydrogel composition | Adhesion ligand | Stiffness* |
| --- | --- | --- | --- |
| | Alginate and gelatin concentration (% w/v) \| Calcium Chloride ($\text{CaCl}_2$, mM) | Gelatin concentration (1% w/v, 3% w/v) | Compressive modulus (kPa) |
| So-L | A1.5G1 \| 100 $\text{CaCl}_2$ | Low | $1.8 \pm 0.2$ |
| So-H | A1.5G3 \| 100 $\text{CaCl}_2$ | High | $2.4 \pm 0.1$ |
| St-L | A3G1 \| 300 $\text{CaCl}_2$ | Low | $6.1 \pm 0.2$ |
| St-H | A3G3 \| 300 $\text{CaCl}_2$ | High | $10.1 \pm 0.5$ |

\*Compressive moduli of selected hydrogels were measured in <sup>25</sup>.

#### 2.2.2 3D breast cancer models: alginate-gelatin hydrogels

MDA-MB 231 cell pellet containing 2×10^6^ cells was re-suspended in 1 mL of alginate-gelatin solution (aq.) using the MICROMAN E viscous pipette (M1000E, Gilson, UK) and ensuring a homogeneous single cell suspension. The cell-suspension was transferred in a sterile 1 mL syringe equipped with a 25G needle. A beaker was filled with sterile CaCl_2_ solution at known concentration (Table 1), the cell suspension was ejected through the nozzle drop-wise, generated hydrogel beads were incubated in the CaCl_2_ solution (aq.) allowing gelation (10 min, RT). Spherical alginate-gelatin hydrogel beads encapsulating cells were recovered using a cell strainer, washed twice with sterile HBS solution, and finally immersed in complete cell culture media and transferred in the incubator.

### 2.3 3D bone models: PCL-based scaffold

3D PCL-based scaffolds (PCL, PCL HA, PCL BaTiO_3_ and PCL SrHA) were manufactured as previously described ^26,27^ by using a 3D-Bioplotter system (EnvisionTEC, Gladbeck; Germany). Briefly, the dispersion phase (10% w/w) of each composite formulation was mechanically mixed with the polymeric phase at room temperature. Subsequently, about 4 g of the raw materials in powder form was introduced into a stainless-steel cartridge and processed as single extruded filament using a 22G nozzle, and according to the printing conditions reported in Table 2. In order to increase pore interconnectivity, porous cylindrical (7 mm diameter) scaffolds were produced with a shifted architecture, made using a laydown pattern of 0/90° and an offset distance equal to half the distance between strands.

**Table 2.** Optimized printing parameters for 3D scaffolds

| Sample ID | Temperature (°C) | Pressure (bar) | Speed (mm/s) | Pre-Flow (s) | Post-Flow (s) |
| --- | --- | --- | --- | --- | --- |
| PCL | 130 | 6 | 0.6 | 0.45 | 0.1 |
| PCL HA | 130 | 6.5 | 0.6 | 0.75 | 0.1 |
| PCL BaTiO<sub>3</sub> | 125 | 5.5 | 0.7 | 0.75 | 0.1 |
| PCL SrHA | 130 | 6.2 | 0.6 | 0.75 | 0.1 |

#### 2.3.1 PCL composite scaffolds: bone scaffold

Composite PCL scaffolds were sterilized as described in our previous study Mancuso *et al*. ^26^. Sterile scaffolds were transferred to a 48 multi-well (MW) plate, then 2×10^5^ Saos-2 cells re-suspended in 50 µL of complete media were gently pipetted on the top of each scaffold. Scaffolds were incubated allowing cell adhesion (30 min, 37 °C, 5% CO_2_), then 400 µL of fresh complete media was added in each well covering the whole scaffold. After seven days of culture (37 °C, 5% CO_2_), the culture media was changed to osteoblast mineralization media (C-27020, PromoCell, UK) to induce mineralization and changed thereafter every four days until the end point (i.e. day 28).

#### 2.3.2 Decellularization of PCL scaffolds: biohybrid bone scaffolds

PCL scaffolds were decellularized via a combination of mechanical and chemical methods ^28,29^, with all steps performed in sterile conditions. After 28 days of culture (end point), scaffolds were washed twice with sterile distilled water and frozen at -80 °C in water overnight. Scaffolds were thawed (2 h, RT), and the freeze-thaw cycle was repeated twice to complete cell lysis. Following the mechanical decellularization step, scaffolds were washed twice with HBS and then incubated with sterile 1 mg/mL DNase (11284932001, Roche) solution diluted in HBS (1 h, 37 °C) to remove any residual nuclear debris. Incubation with 0.05% (w/v) SDS solution in HBS (15 min, RT) to remove any further remaining debris. Before further cell culture experiments, scaffolds were washed (n = 3) with HBS (5 min, RT) to remove any residual reagents.

### 2.4 Alginate-gelatin hydrogels and PCL scaffold: breast-to-bone model

#### 2.4.1 Indirect migration model

The *indirect migration model* was obtained by seeding pre-conditioned breast cancer cells onto biohybrid bone scaffolds. Prior this step, MDA-MB 231 cells were cultured in alginate-gelatin hydrogels (Table 2) for 7 days, allowing conditioning in distinctive breast tumor microenvironments (Figure 1A). After the pre-conditioning step, MDA-MB 231 cells were retrieved from alginate-gelatin beads using the dissolution buffer. As control, MDA-MB 231 (non-conditioned) and MDA-IV cells (non-conditioned) were cultured on standard tissue culture plastic (TCP) wells, and detached with trypsin (T3924, Sigma-Aldrich, UK) following standard protocols (3 min, 37 °C). All recovered cells were centrifuged at 600 g and re-suspended in complete media, then seeded directly on biohybrid scaffold at a seeding density of 1×10^5^/scaffold, and on TCP wells at density of 2 × 10^4^ cells/cm^2^ as additional control group (Figure 1A). For all the mentioned conditions, experiments were performed in complete cell culture media -/+ 5 ng/mL TGF-β (100-21, Peprotech). For all the conditions, cells were cultured (37 °C, 5% CO_2_) for 7 days in complete DMEM medium, changing the media every 2 days.

**Figure 1:**
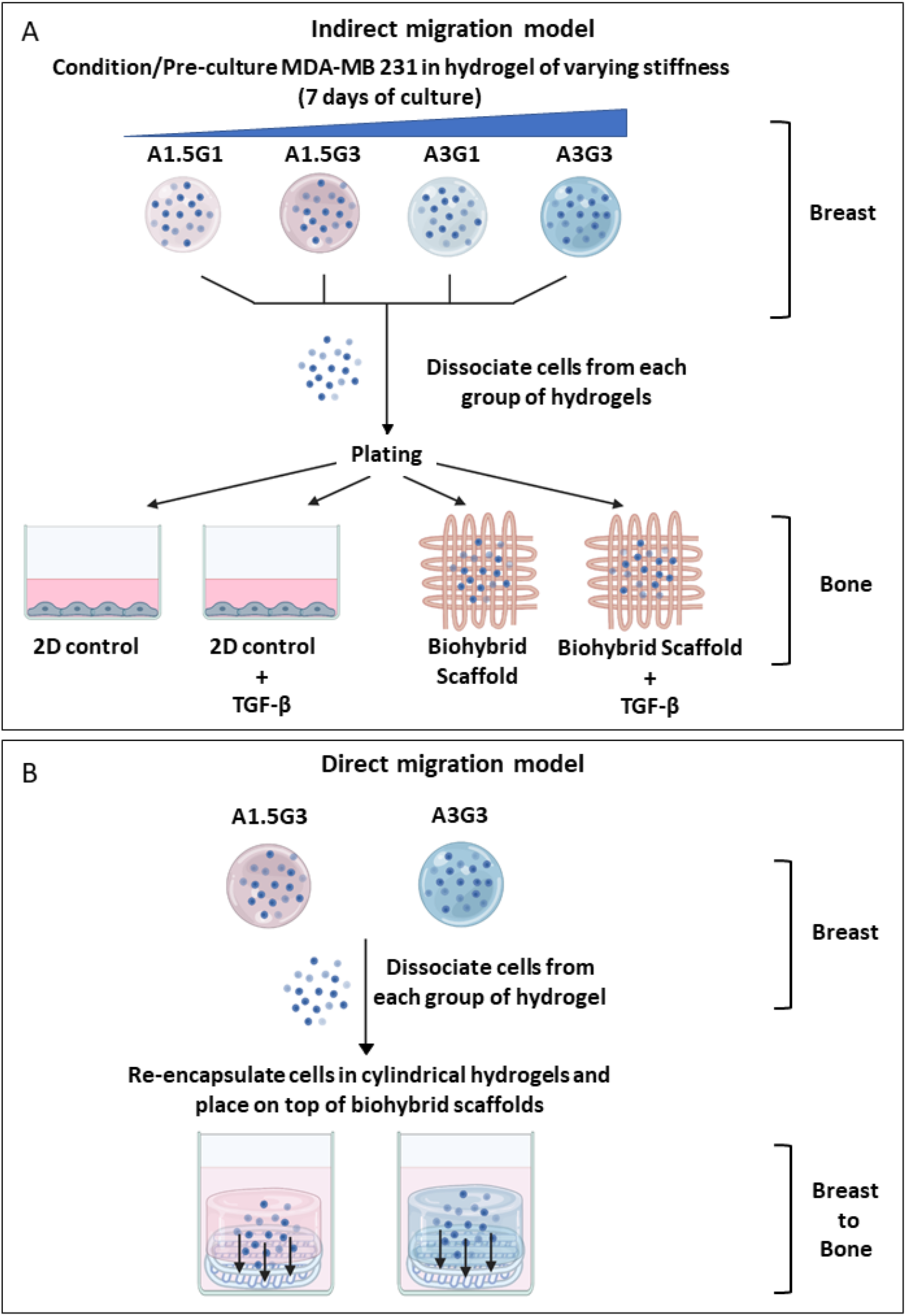
Schematic layout of experimental plan for *indirect* (A) and *direct migration* (B) models.

#### 2.4.2 Direct migration model

The *direct migration model* was designed to mimic the migration of breast cancer cells from primary (breast) to the secondary site (bone), aka from hydrogels to biohybrid scaffolds (Figure 1B). Briefly, cells were retrieved from hydrogels after 7 days using the dissociation buffer and re-encapsulated in the same hydrogel at a density of 2×10^6^ cells/mL. 200 µL of cell-hydrogel suspension was gently pipetted on top of the decellularized PCL scaffold in a 48 MW plate and incubated (15 min, 4 °C) to allow physical gelation of gelatin and obtain cylindrical shaped cell-laden hydrogels. Then 200 µL of CaCl_2_ (aq.) was added to the well allowing ionic alginate crosslinking (10 min, RT). The combined scaffold was washed with HBS (n = 3) and then supplemented with complete DMEM. Cells were incubated up to 7 days and cell culture media was replaced every day.

### 2.5 Cell proliferation assay

Alamar blue assay (Deep Blue Cell Viability™ Kit, 424701, Biolegend) was used to analyze proliferation of MDA MB 231 and MDA-IV cells in PCL scaffolds at day 1, 3 and 7. Briefly, cell culture media was gently removed from each well, 400 µL of deep blue solution (10% v/v Deep blue viability reagent in complete cell culture media) was added to each well and incubated for 2 hours (37 °C, 5% CO_2_). Then, 200 µL of cell culture media was taken from each well, transferred to a 96 well plate, and immediately measured with Synergy-2 (Biotek) plate reader (Ex 530-570 nm / Em 590-620 nm). The measurements are reported as mean ± standard deviation (SD) of duplicates samples (n = 2) and N = 3 independent experiments.

### 2.6 Adhesion/ Cell spreading assay

8-well chambers slides (80826, Ibidi) were coated with collagen according to manufacturer’s instructions. Briefly, each chamber was coated with 35 µg/mL Collagen type I (sterile, 50201, Ibidi) diluted in 17.5 mM acetic acid (aq.) and incubated (1 h, RT). Collagen solution was slowly removed and chambers were washed with sterile PBS (n = 1). Similarly, fibronectin (F2006, Sigma) was diluted in sterile PBS and used at a concentration of 20 µg/mL (1 h, RT). After incubation, fibronectin solution was removed and chambers were washed with sterile PBS (n = 1). After washing, both collagen and fibronectin coated chambers were left to air dry in sterile conditions (30 min, RT).

Cells pre-conditioned in selected hydrogels (7 days) were recovered using the dissociation buffer and seeded on uncoated (control), collagen or fibronectin coated surfaces at a density of 1×10^4^ cells/cm^2^. After 45 minutes of incubation (37 °C, 5% CO_2_), cells were fixed with 4% paraformaldehyde (PFA) and incubated (30 min, RT) with 1 µg/mL DAPI in PBS (D954, Sigma-Aldrich, UK) and a 1: 50 phalloidin Alexa-568 in methanol (A12380, Invitrogen). Approximately n = 200 cells were imaged per condition (Section 2.12.1).

### 2.7 Scratch assay

MDA-MB 231 cells pre-conditioned in hydrogels for 7 days (Table 2) were recovered as previously described, seeded in 6-well plate at a density of 4×10^5^ cells/well and transferred to the incubator (37 °C, 5% CO_2_) allowing cell adhesion (24 h). A scratch was then performed in each well using a sterile 200 µL tip, each well was washed with cell culture media to remove any cellular debris. Cells were supplemented with low serum media (1% v/v FBS in DMEM with 1% v/v L-glutamine and 1% v/v PenStrep) to reduce cell proliferation ^30,31^. Area of the scratch covered was measured at day 1 and day 2 (Section 2.12.2) The experiments were performed in duplicates (n = 2) and in N = 3 independent experiments and the data presented as mean ± SD for each sample.

### 2.8 Invasion assay

Silicon inserts (80406, Ibidi) were cut in 4×4 mm pieces placed at the center of each well (8-well chambers, Ibidi) to create a cell free space (Figure S1A). Collagen (50201, Ibidi) precursor mix was prepared following the supplier’s instruction: collagen was diluted to a final concentration of 1 mg/mL in 10× DMEM (D2429, Sigma-Aldrich) and sterile water, the pH was adjusted to 7.2-7.4 using sterile 1 M NaOH (aq.) (12963614, Fischer scientific) and 7.5% w/v NaHCO_3_ solution (S8761, Sigma-Aldrich). MDA-MB 231 cells were gently re-suspended in the collagen pre-gel solution at a density of 7.5×10^5^ cells/mL. Immediately, 200 µL of cells-collagen mix was pipetted outside the silicon insert and incubated allowing collagen gelation (30 min, 37 °C, 5% CO_2_). The central insert was removed and 50 µL of 1 mg/mL collagen solution were pipetted to fill the space with cell-free collagen hydrogel. The whole setup was further incubated to complete gelation (20 min, 37 °C, 5% CO_2_).

After gelation, encapsulated cells were stained with Cytopainter red (ab138893, Abcam). Briefly, cells were incubated with 1× dye diluted in cell culture media for 1 hour in the incubator and washed twice with 1× PBS followed by addition of cell culture media only. The invasion area was measured at day 3 (Section 2.12.3). The experiments were performed in duplicates (n = 2) and in N = 3 independent experiments and the data presented as mean ± SD for each sample.

### 2.10 PThrP expression

PThrP expression was detected from cell lysates at day 7 using the Human PTHrP Elisa kit (E-EL-H1478, Elabscience). MDA-MB 231 cells were first trypsinized from the either scaffolds or well plates (5 min, 37 °C), then recovered using fresh media. Cells were centrifuged (3 min, 500g), supernatant removed and the pellet was incubated (5 min, 4 °C) in 200 µL of lysis buffer composed of 1× RIPA buffer (ab156034, Abcam), complete mini protease inhibitor cocktail tablet (11836170001, Roche) and 1 µM phenylmethylsulfonyl fluoride (PMSF). After 5 minutes on ice, the cell lysate was centrifuged (10 min, 4 °C, 2000g) and supernatant collected for further ELISA analysis. Sandwich ELISA was performed as per supplier’s manual. The blank OD values (lysate buffer only) obtained at 450 nm were subtracted from sample and standard’s OD values and pg/mL values of samples were calculated using the standard calibration curve. BCA protein assay (23228, Thermo Scientific) was used to measure the lysate concentration for each sample. Finally, the obtained PThrP concentrations were further normalized by their respective lysate concentration, obtaining PTHrP pg/mg of protein values. The measurements were carried in duplicates (n = 2) for each experiment and values are plotted as mean ± SD of N = 3 independent experiments.

### 2.11 IL-6 release quantification

For each sample, fresh media was changed on day 6 and collected on day 7 (i.e. 24 hours later). Briefly, media was centrifuged (10 min, RT, 1000g) and the supernatant collected and used to detect IL-6 release. ELISA (human IL-6 Kit, 550799, BD OptEIA™) was performed according to supplier’s instruction, with fresh complete media used as blank. The amount of IL-6 (pg/mL) was determined from the standard calibration curve of known standard IL-6 concentration. Obtained concentrations were normalized against their respective cell proliferation data (Section 2.5) measured at day 7. The measurements were carried in duplicates (n = 2) for each experiment and values are plotted as mean ± SD of N = 3 independent experiments.

### 2.12 Image acquisition and analysis

Images were acquired using the fluorescent inverted microscope (Leica DMI6000, Leica Microsystems, UK) coupled with a 5.5 Neo sCMOS camera (Andor, UK), and equipped with: 2× objective (PLAN 2.5×/0.07, Leica), dry 10× objective (PL 10×/0.3 PH1, Leica), dry 20× objective (PL 20×/0.5 PH2, Leica), dry 63× objective (PL 63×/0.9 PH2, Leica), and filter cubes (A4, I3 and N2.1). The µManager software (v.1.46, Vale Lab, UCSF, USA) was used to control both microscope and camera, as well as to capture images.

#### 2.12.1 Adhesion assay

Cells were imaged with the 20× dry objective with filter cubes A4 and N2.1. Images were analyzed with ImageJ (v1.49p) for object identification and measure the cell spread area. Nearly 200 cells were analyzed per condition, and individual cell area was plotted for each condition.

#### 2.12.2 Scratch assay

Brightfield images of the scratch were taken using the 10× dry objective at different time points (day 0, day 1 and day 2). The area of scratch invaded by cells was calculated using ImageJ by measuring the combined cellular area in the scratch over time.

#### 2.12.3 3D collagen invasion assay

Each well was imaged using the 2× objective and N2.1 filter cube, at different time points (day 0 and day 3). In order to measure invasion on the overall volume, z-stacks (with z-step of 50 µm to get single cell resolution) were acquired, and the maximum projection of each sample was analyzed with ImageJ. For the analysis, maximum projections were converted to binary, and the total area stained in the acellular collagen hydrogel at day 3 was measured for each condition. Images were acquired at day 0 and day 3.

### 2.13 Statistical analysis

Significance for cell adhesion assay was done by non-parametric one-way ANOVA with Dunn’s post-hoc multiple comparison test. Both migration and invasion assay were analyzed with one-way analysis of variance (ANOVA) followed by Tukey’s post-hoc multiple comparison test. For PTHrP and IL-6 analysis significance among conditions were analyzed by Two-way ANOVA followed by Tukey’s post-hoc test. All significance test and plotting of graphs was done by using GraphPad prism v9.1.0. P-values were set at four different significance levels: *p ≤ 0.05, **p ≤ 0.01, ***p ≤ 0.001, ****p ≤ 0.0001.

## 3. Results and Discussion

### 3.1 Conditioning MDA-MB 231 with high stiffness leads to increased migratory and invasive phenotype

MDA-MB 231 are well known for their invasive potential and are commonly used as a model to study bone metastasis ^32,33^. In this study, MDA-MB 231 were selected and used to understand how ECM properties could impact on various aspects of their metastatic potential. We examined effect of stiffness and composition of hydrogels on MDA-MB 231’s invasive and migratory phenotypes using three experiments: adhesion ability (2D cell spread assay, paragraph 2.6), migration ability (2D scratch assay, paragraph 2.7) and invasion (3D collagen invasion assay, paragraph 2.9). In all experiments, MDA-MB 231 cells were allowed to adapt to the microenvironment (i.e. hydrogels) for 7 days: cells were encapsulated in alginate-gelatin hydrogels with known stiffness and composition (Table 1).

Adhesion assay examined cellular adhesion and extent of membrane protrusion, both of which precede migratory and invasive phenotype. Substrate adhesion was examined by plating pre-conditioned MDA-MB 231 cells on non-coated, collagen and fibronectin coated plates (Figure 2A-D). The number of cells that attached to the substrate (adhesion) didn’t vary significantly among groups of pre-conditioned cells in all the substrates tested. Also, individual cell spread area of pre-conditioned cells exhibited no differences on non-coated and collagen coated plates. On the contrary, cells preconditioned in St-H hydrogels (i.e. high stiffness, high gelatin) showed increased spread area in fibronectin coated plates (Figure 2A, 2D). In this model, both high stiffness and composition of hydrogels are responsible for changes in cells’ adhesion capacity.

**Figure 2.**
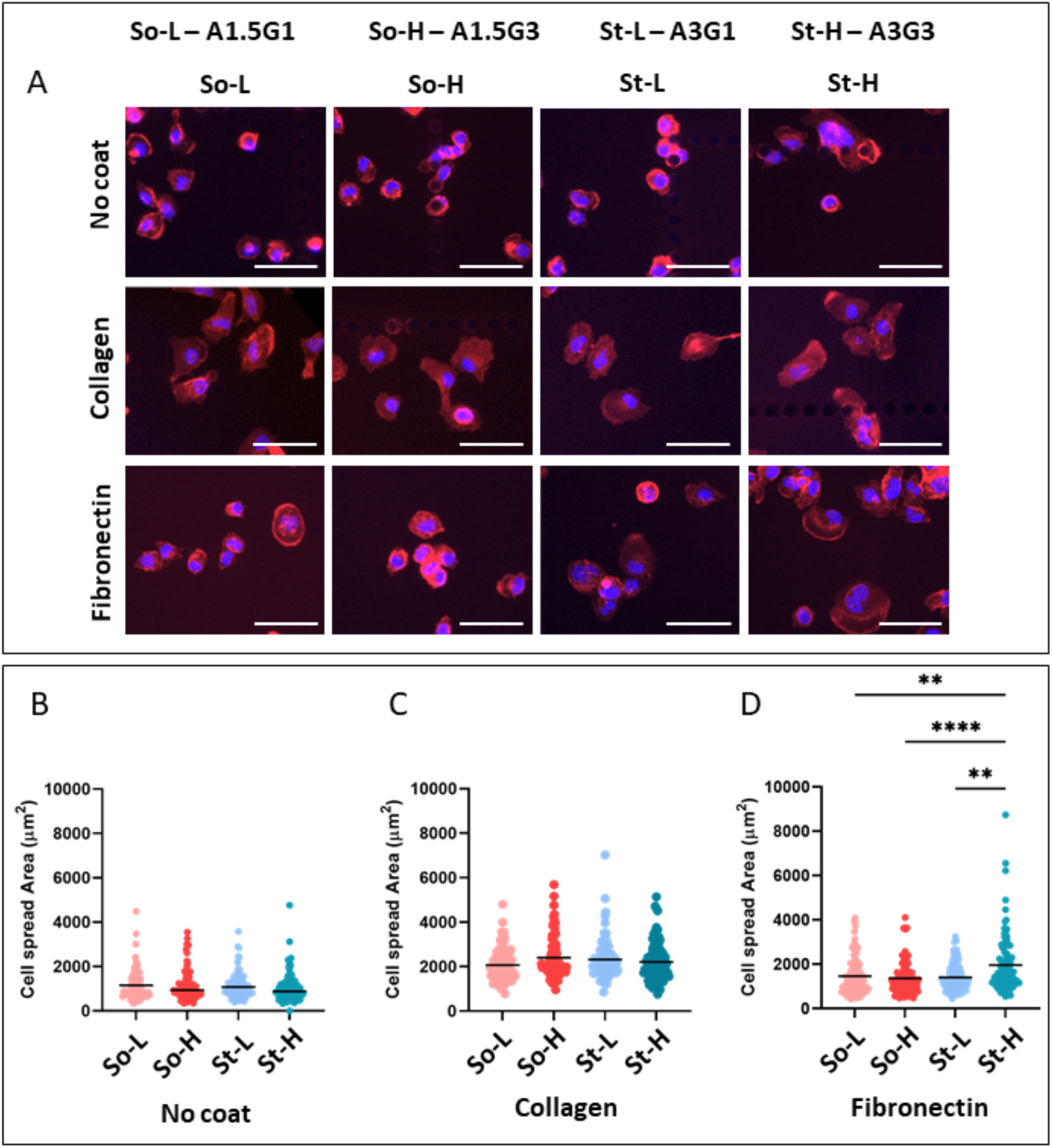
Cell spread area assessed on non-coated, collagen and fibronectin covered plates. **A)** Immunofluorescent images of MDA-MB 231 cells stained with DAPI (nuclei, blue) and Phalloidin (F-Actin, red) conditioned in the selected four alginate-gelatin hydrogels on non-coated, collagen-coated or fibronectin-coated surfaces (Scale bars: 100 µm). Dot-plot representations of individual cell area and mean of at least n=200 cells of: **B)** non-coated, **C)**, Collagen-coated, and **D)** and Fibronectin-coated. P-values represented as *p ≤ 0.05, **p ≤ 0.01, ***p ≤ 0.001, ****p ≤ 0.0001).

Parallel to previous findings, MDA-MB 231 cells pre-conditioned in stiff hydrogels (compressive modulus > 6 kPa) covered the scratch area significantly faster at both days than the ones from softer hydrogels (compressive modulus < 3 kPa) as shown in (Figure 3A-C). Cells conditioned in So-L vs St-L hydrogels showed 30% increase (p < 0.01) and So-H vs St-H showed 40% increase in covering the scratch area (p < 0.0001) at Day 2 (Figure 3C). Within high stiffness groups, cells cultured in St-H showed increased migration capacity than St-L (p ≤ 0.001, Figure 3C). These results suggest a primary role of stiffness in this phenotype with gelatin content affecting cell migration capacity only when coupled with high stiffness.

**Figure 3.**
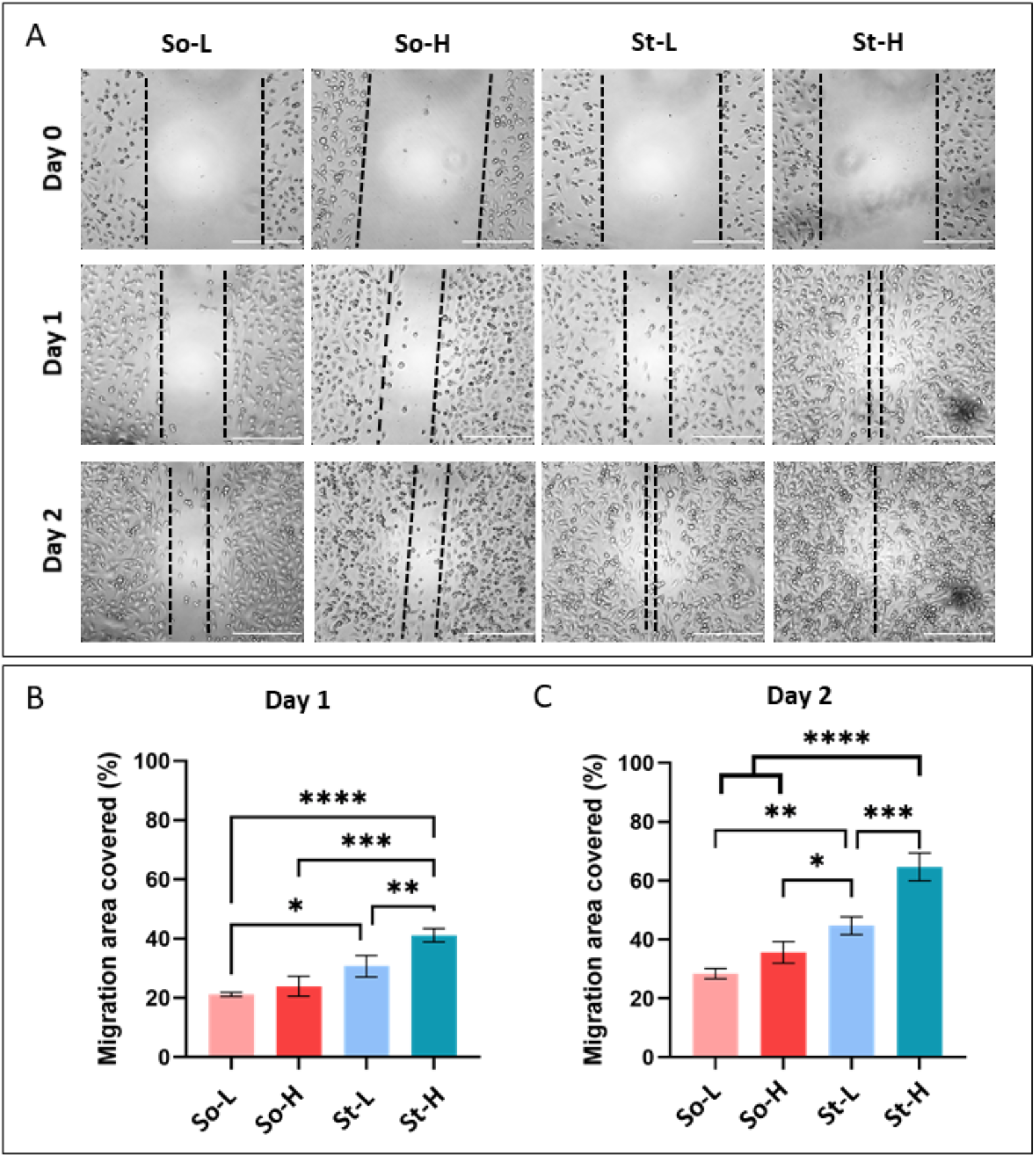
Scratch assay performed on stiffness conditioned cells: **A)** Brightfield images of the scratch assay performed on cells conditioned in four hydrogels at day 0, day 1 and day 2 (Scale bars = 500 µm). Dashed black lines are used as visual representation and to help identifying cell migration over time in the different conditions tested. Graphical representation of scratch area (measured as µm^2^) covered by migratory cells is represented as % in comparison to blank area measured at day 0 for **B)** day 1 and **C)** day 2. The data is represented as mean ± SD of n=2 wells, N=3 independent experiments. P-values represented as *p ≤ 0.05, **p ≤ 0.01, ***p ≤ 0.001, ****p ≤ 0.0001).

Invasive potential in 3D was quantified measuring the cells ability to invade pristine collagen hydrogels. Again, MDA-MB 231 cells pre-conditioned in selected alginate-gelatin hydrogels were resuspended in collagen at known concentration and migration towards acellular collagen hydrogels was observed over time. To measure invasion, MDA-MB 231 cells were stained with cytopainter red, as this staining is retained only by cells encapsulated in the gel at day 0. In this way, daughter cells proliferated in collagen were removed from the counting, allowing to distinguish invasion from cellular proliferation (Figure 4A, Figure S1B). After three days, cells from stiffer hydrogels (St-L and St-H) invaded the area 1.5 times more than the ones from softer hydrogels (So-L and So-H, Figure 4B). We did not observe any correlation between the amount of gelatin in hydrogels and cells’ invasive potential (Figure 4), and no statistical difference was observed when comparing hydrogels with similar stiffness and increased gelatin content (So-L vs So-H, St-L vs St-H). On the contrary, and as expected, cells from stiffer hydrogels have higher invasive potential compared to cells from softer hydrogels (So-L vs St-L, p ≤ 0.01; So-H vs St-H, p ≤ 0.05).

**Figure 4.**
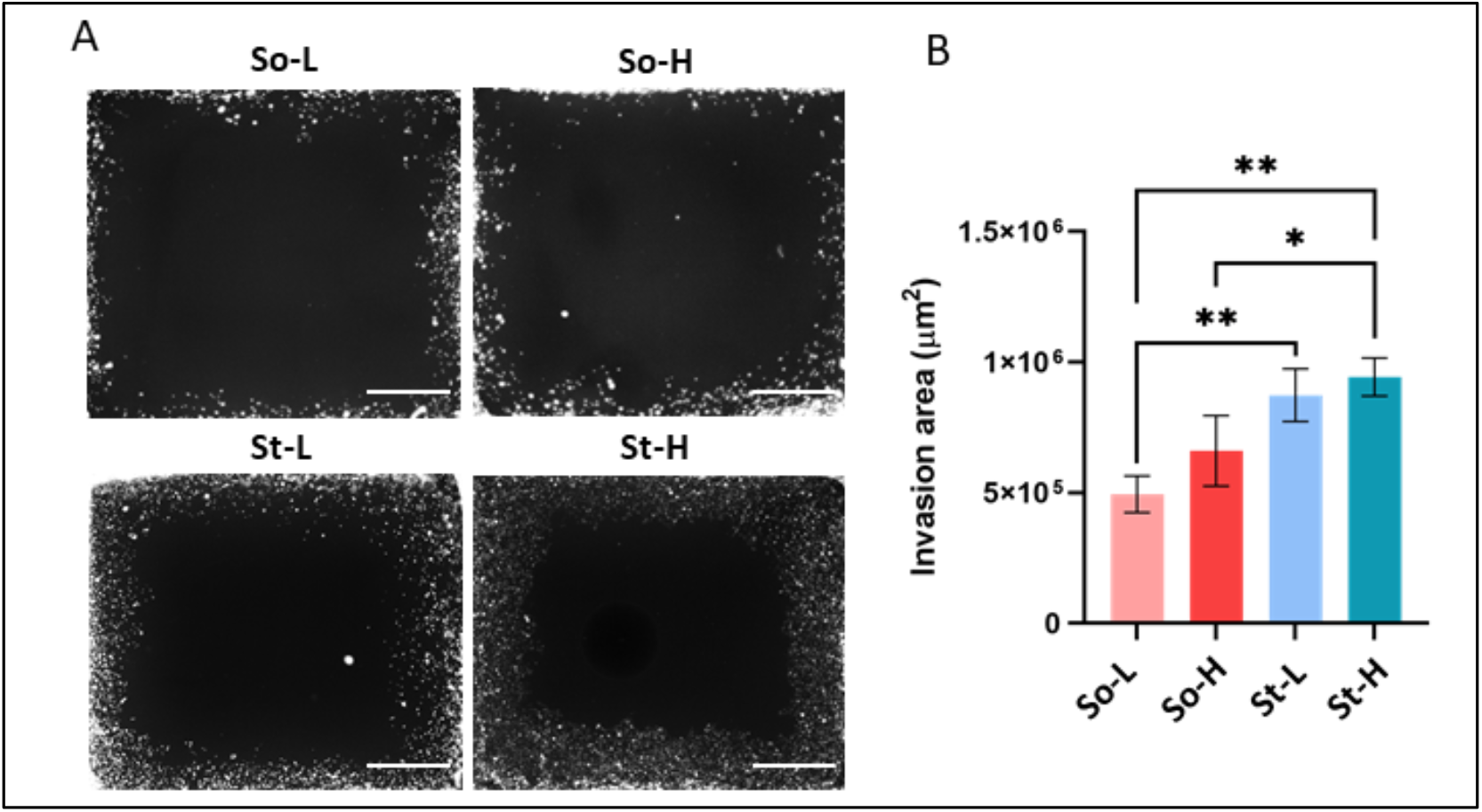
3D collagen invasion assay of stiffness conditioned cells. **A)** Immunofluorescence images of Cytopainter red stained MDA-MB 231 cells encapsulated in outer collagen layer at day 0 and day 3. **B)** Cropped images of the acellular collagen hydrogel area showing invasion of MDA-MB 231 pre-conditioned in alginate-gelatin hydrogels at day 3 to showcase impact of alginate-gelatin hydrogels on cell invasive potential (Scale bars = 1000 µm). **C)** Binary images of cropped area obtained after post-processing were used to calculate the invaded area (µm^2^) and represented as mean ± SD of n = 1 and N = 3 independent experiments. P-values represented as *p ≤ 0.05, **p ≤ 0.01, ***p ≤ 0.001, ****p ≤ 0.0001).

The results suggest that pre-conditioning cells in environments with different stiffnesses plays a major role in directing MDA-MB 231 migratory and invasive phenotypes, whereas this correlation was not evidenced at different adhesion ligand concentrations. While this study only elucidates effects of gelatin-related adhesion motifs, inclusion of other ECM components like fibronectin, laminin, and hyaluronic acid could better illustrate contributions of ECM composition on migration and invasion capacity ^3^.

### 3.2 Decellularized PCL scaffolds to mimic bone ECM

PCL scaffolds are known to retain ECM deposited by cells cultured on them after decellularization ^28,29^. Here, we compared osteogenic potential of composite PCL scaffolds and evaluated their ECM retaining ability after decellularization (Figure 5A). We selected four PCL-composite scaffolds (i.e. PCL, PCL HA, PCL SrHA and PCL BaTiO_3_) to mimic bone tissue properties ^26,27^. All the scaffolds were 3D printed with the same 3D architecture and characterized in terms of composition, strand dimension and mechanical properties ^26,27^. PCL-composite scaffolds were cultured with Saos-2 up to 28 days, allowing deposition of bone-ECM. Decellularization of scaffolds was performed to assess: A) the capacity to retain ECM-deposited, and B) the feasibility to remove cells from scaffolds in sterile condition and allowing for their use as metastatic site in other experiments (Figure 5A). SaOs-2 cells were chosen as human osteoblast model because of their resemblance to osteoblastic properties especially matrix production and calcium deposition under mineralizing conditions ^34–36^. Osteoblasts maturation was quantified in all PCL-based scaffolds up to 28 days using: cell proliferation, ALP activity, calcium deposition (Alizarin stain) and ECM deposition (collagen and osteocalcin IF stain) ^37^. Proliferation data suggests higher compatibility of composite PCL scaffolds (PCL HA, PCL SrHA, PCL BaTiO_3_) when compared to pristine PCL ones (Figure S2A). PCL BaTiO_3_ scaffolds were found to induce highest amount of ALP in Saos-2 cells when compared to other scaffolds (Figure S2B); moreover, both PCL HA and PCL BaTiO_3_ scaffolds lead to increased calcium deposition by Saos-2 cells (alizarin stain, Figure S3). IF stain confirmed deposition of both collagen and osteocalcin, suggesting comparable deposition of both ECM proteins by SaOs-2 on all scaffolds tested by day 28 (Figure S4).

**Figure 5.**
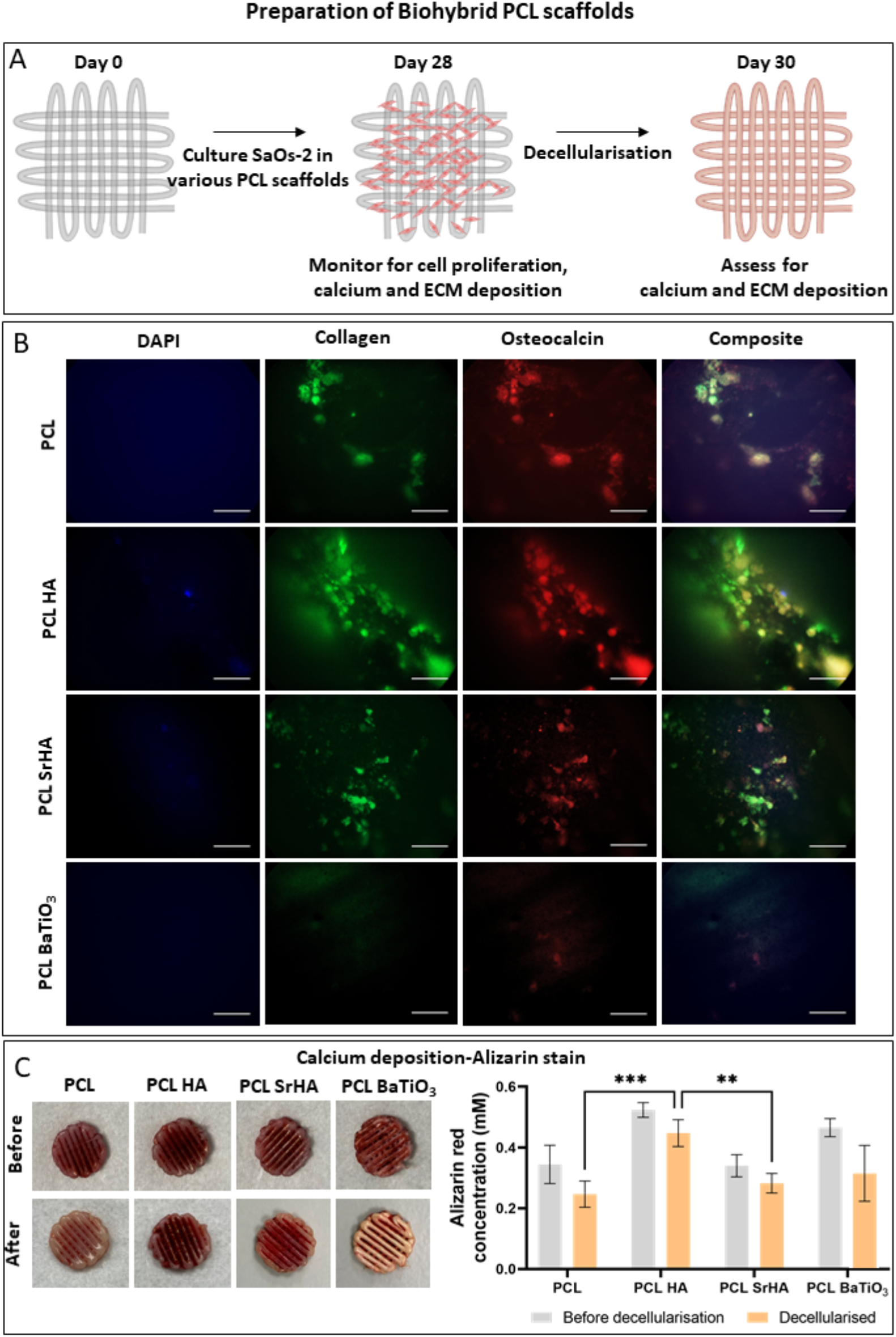
Characterization of decellularized composite PCL scaffolds to mimic bone ECM. **A)** Schematic representation of preparation of decellularized scaffolds and their characterization. **B)** Immunofluorescent images of nucleus (blue), collagen (green) and osteocalcin (red) of stained PCL-based scaffolds after decellularization steps (Scale bars: 50 µm). **C)** Images of PCL-based scaffolds stained with Alizarin stain before and after decellularization and Alizarin stain concentration (mM) from scaffolds before and after decellularization. Values are represented as mean ± SD of N=3 independent experiments. P-values represented as *p ≤ 0.05, **p ≤ 0.01, ***p ≤ 0.001, ****p ≤ 0.0001).

The ability to use decellularized PCL scaffolds and to create sterile *biohybrid PCL scaffolds* (i.e. composite bone-mimicking scaffolds enriched in bone mineralized matrix) was assessed as requirement for further use of biohybrid scaffolds in other in vitro experiments. In fact, *biohybrid PCL scaffolds* could act as a more physiologically relevant in vitro secondary site for breast cancer. The decellularization steps were optimized for all PCL-based scaffolds (Supplementary Information, Figure S5), and the incubation with 1 mg/mL of DNase to lyse nuclear debris after mechanical disruption of cells was found critical to completely remove any residual of Saos-2 cells (Figure S5, 5B). The capability to retain deposited ECM was tested quantifying the amount of collagen and osteocalcin in decellularized scaffolds (IF staining, Figure 5B). Pristine PCL, PCL HA and PCL SrHA scaffolds retained both collagen and osteocalcin; whereas, PCL BaTiO_3_ showed minimal ECM proteins with negligible staining detected (Figure 5B). Calcium deposition after decellularization was measured using Alizarin stain, observing a minimal reduction of calcium in all scaffolds tested. Of note, the highest calcium amount detected was in PCL HA scaffolds (Figure 5C). Pristine PCL scaffolds and PCL BaTiO_3_ scaffolds lost around 30% of previously quantified calcium deposition, which is possible due to the decellularization process. Both PCL HA and PCL SrHA scaffolds were instead able to retain more calcium (about 10% reduction with respect to Saos-2 colonized scaffolds). Of note, Alizarin stain also recognizes intracellular calcium, hence the reduction observed after decellularization could be attributed also to loss of intracellular calcium. From the results, PCL HA scaffold was selected as secondary metastatic scaffold as the one able to retain the highest amount of calcium and deposited ECM, which will from now on referred to as biohybrid PCL HA scaffold.

### 3.3 *Indirect migration* of MDA-MB 231 and MDA-IV cells in biohybrid scaffolds

We have shown that pre-conditioning MDA-MB 231 cells in different alginate-gelatin hydrogels of varying stiffness and adhesion motifs concentration could impact on invasive potential and migratory phenotypes of breast cancer cells (Figure 1-3). Further in this study, we used two models to study the impact of pre-conditioning on breast-to-bone metastasis. We firstly investigated the *indirect migration model*, in which pre-conditioned MDA-MB 231 cells were seeded onto biohybrid PCL HA scaffold. This model was designed to study the response of breast cancer cells to bone ECM, irrespective of their bone localizing potential. As a control, non-conditioned MDA-MB 231 and MDA-IV (i.e. bone homing variant of MDA-MB 231) were cultured on the same biohybrid PCL HA scaffold. Expression of osteolytic factor (i.e. PTHrP, IL-6) and cell proliferation were examined up to 7 days of culture. To further investigate the role of bone microenvironment growth factors released in response to increased osteoclast activity in bone metastasis, 5 ng/mL TGF-β was supplemented as additional variable of the model (Figure 6).

**Figure 6.**
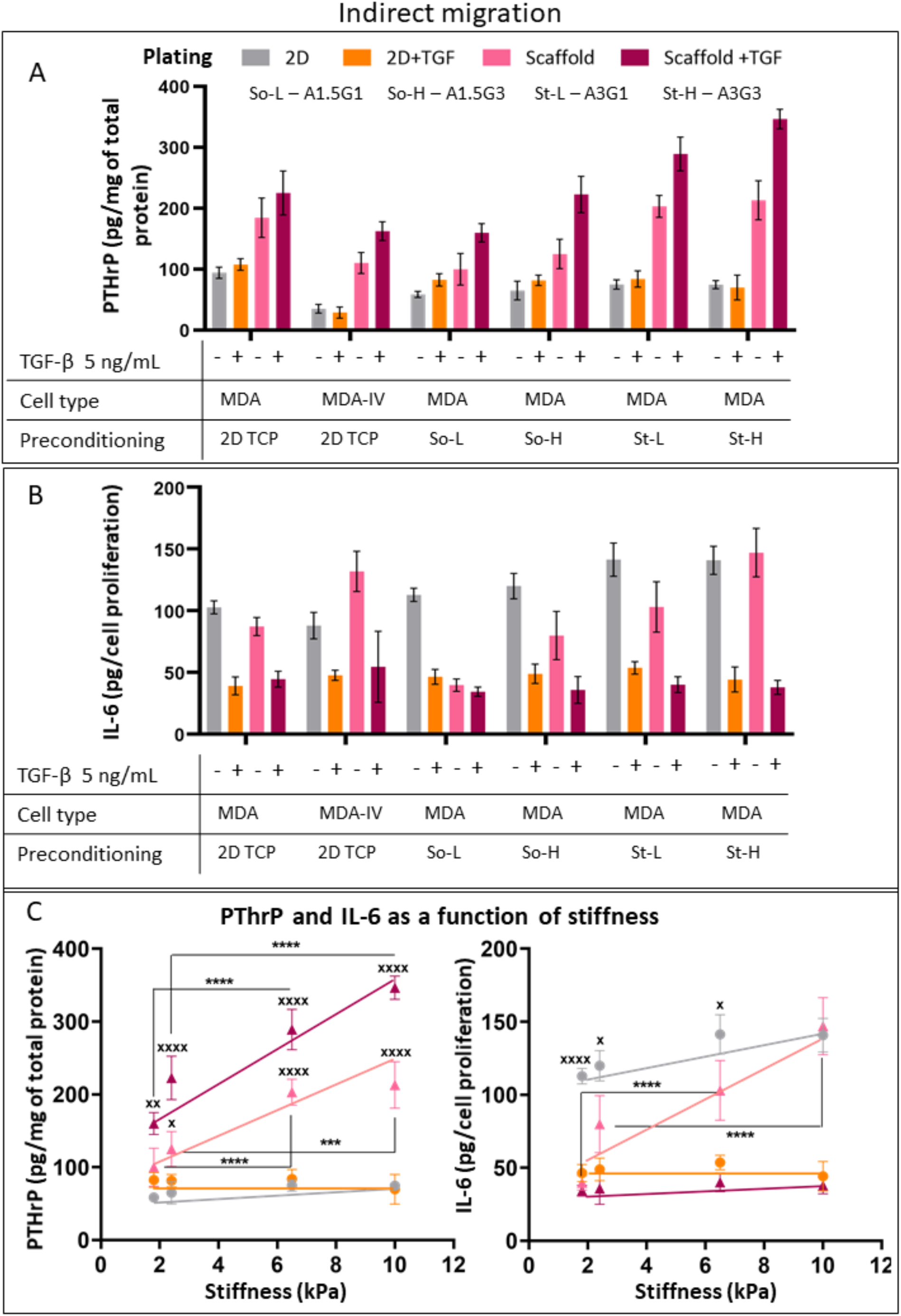
PTHrP expression and IL-6 release in *indirect migration model*. **(A)** PTHrP expression and (B) IL-6 release quantified in MDA-MB 231 (referred to as MDA) and bone homing MDA-IV cells plated on either TCP plates (**Grey** -without TGF-β, and **Orange** -with TGF-β) or biohybrid PCL HA scaffolds (**Pink -**without TGF-β and **Purple** -with TGF-β). MDA-MB 231 cells were pre-conditioned in TCP plates, or four groups of alginate-gelatin hydrogels as mentioned below each graph. The values are represented as mean and SD of n=2 scaffolds and N=3 independent experiments. Same data of PTHrP expression and IL-6 release in MDA-MB 231represented as a function of stiffness values (kPa) of hydrogels in which they were pre-conditioned **(C)**. P-values represented as *p ≤ 0.05, **p ≤ 0.01, ***p ≤ 0.001, ****p ≤ 0.0001).

PTHrP expression as measured by ELISA at day 7 of culture showed overall increased expression in cells plated on biohybrid PCL HA scaffolds compared to controls (i.e. 2D TCP plates), confirming that exposure to bone specific ECM is essential to induce PTHrP expression (Figure 6A), which is a first key step in bone metastasis cycle. Interestingly MDA-MB 231 cells cultured on biohybrid PCL HA scaffolds showed proportional expression of PTHrP with stiffness of hydrogels (preconditioning step); two-fold increase was observed when comparing stiffer conditioned groups than softer (p < 0.0001 for So-L vs St-L, p < 0.001 for So-H vs St-H) (Figure 6A, 6C). The same trend (proportional PTHrP expression with stiffness) was observed in the presence of TGF-β (p < 0.0001 for So-L vs St-L and So-H vs St-H). Overall, the presence of TGF-β lead to an increased PTHrP expression in biohybrid PCL HA scaffolds (Figure 6C). Of note, no correlation of PTHrP expression with stiffness was observed when MDA-MB 231 were cultured on TCP and regardless the presence of TGF-β in the cell culture media. This trend suggests the active role of A) the primary site and cellular adaptation to the stiffness of the microenvironment and B) the composition of the secondary site in promoting osteolytic activity.

IL-6 release was assessed in the same way, we found that in absence of TGF-β, IL-6 release increased two-fold when cells were pre-conditioned in high stiffness hydrogels and seeded on the biohybrid scaffolds but remained constant in TCP controls (Figure 6B). However, unlike PTHrP, IL-6 released by cells plated on scaffolds was less compared to that of TCP except for cells pre-conditioned in St-H hydrogels that matched the 2D control (Figure 6B, 6C). Surprisingly, with TGF-β supplementation IL-6 release was reduced in all the tested conditions with values remaining constant in both conditioned and non-conditioned cells (Figure 6B, 6C).

No significant difference in MDA-MB 231 proliferation (day 3, day 7) in biohybrid PCL HA scaffolds was observed between conditioned and non-conditioned cells (Figure S6A). However, the presence of TGF-β induced an increase in proliferation ^26^, found proportional with alginate-gelatin stiffness used to pre-condition cells. Of note, MDA-IV cells growth was found similar to MDA-MB 231 pre-conditioned in the stiffer and high gelatin content hydrogel (St-H, Figure S6B).

In summary, MDA-MB 231 cells express less PTHrP when cultured on TCP plates but show an increased expression in response to high stiffness conditioning from primary tumour and to exposure to bone ECM (secondary site). These results support the need to use models representing better the characteristics of tissues (i.e. 3D/scaffolds) rather than conventional models (i.e. 2D/TCP surfaces) as they mimic microenvironment of the pathology. On the other hand, IL-6 is released at higher concentration by MDA-MB 231 on TCP plates, with release correlated and fine-tuned by stiffness of the pre-conditioning microenvironment when exposed to bone ECM.

The addition of TGF-β on PTHrP and IL-6 release aligns with previous studies, where a clear relationship of increased PTHrP expression induced by TGF-β in breast-to-bone metastasis is reported ^20,38^; and no correlation with IL-6 expression in the same context. Studies done in other model systems report complex and mixed crosstalk between TGF-β and IL-6. In intestinal epithelial cells, TGF-β dampens IL-6 signaling but reports no direct effect on its expression ^39^. In biliary tract cancer, they work synergistically to induce EMT and chemotherapy resistance_40_. Both TGF-β ^41^ and IL-6 ^42^ have pleiotropic effects which makes their interaction complex, hence further investigation is needed to draw solid conclusions about their interaction within this model.

Results show that the *indirect migration model* can recreate initial steps of bone metastatic cycle, and in particular the release of osteolytic factors by cancer cells in response to bone ECM. Especially, high stiffness of the in vitro primary tumour is linked to increase in both PTHrP and IL-6 expression when cancer cells are exposed to bone ECM. Of note, colonization is a later stage of the metastatic cycle and relies on complex interactions with other components such as cell types (e.g. osteoblasts, osteoclasts) and tissue vascularization. Hence, an in-depth understanding of metastatic onsets could be achieved by including relevant bone cells in the model, enabling to capture significant and relevant in vivo interactions. This could further help in better understanding specific biological processes to support animal and clinical studies.

### 3.4 *Direct migration* of MDA-MB 231 and MDA-IV from hydrogels to biohybrid scaffolds

The *direct migration model* was designed to study the localization / migration of cells into the bone ECM by placing alginate-gelatin hydrogels containing breast cancer cells (primary site) on the top of *biohybrid PCL scaffolds* (secondary site). Two hydrogels varying in stiffness but with similar gelatin concentration were selected (i.e. So-H and St-H), as stiffness was found to be the primary parameter dictating MDA-MB 231 response in the *indirect migration model* (Figure 6). After pre-conditioning in So-H and St-H, both MDA-MB 231 and MDA-IV cells were re-encapsulated in the same hydrogel and cultured on top of *biohybrid PCL scaffolds* to study migration to the secondary site (as shown in Figure 1).

To assess migration, *biohybrid PCL scaffolds* were separated from hydrogels at the endpoint (day 7) and the number of viable cells migrated into them was determined using the Deep blue viability assay. Results were observed to be similar between MDA-MB 231 and bone homing MDA-IV cells, where higher number of cells migrated from softer hydrogel than stiff hydrogel (So-H > St-H, Figure S7). These results apparently do not support what is observed in migration and invasion assays (Section 3.1), where a correlation of the latter is observed with pre-conditioning in stiffer hydrogels. Literature supports both these observations, where invasion of cancer cell is high within softer alginate gels ^43^, but when cultured in a separate collagen gel, cancer cells pre-conditioned in stiff matrix are found to invade more than the ones pre-conditioned in soft matrix ^24^. The possible explanation could be that stiff and dense matrix can constrict migration but when such conditioned cells reach surrounding soft/less dense matrix, they have higher potential to invade.

Interestingly, PTHrP expression in migrated MDA-MB 231 cells pre-conditioned in St-H hydrogels was found 1.4-fold higher than So-H hydrogels (p < 0.05) (Figure 7A). Similarly, IL-6 release was measured to be 8-fold higher in cells pre-conditioned in stiffer hydrogels (St-H vs So-H, p<0.0001) (Figure 7B). This suggests that while less cells migrated from high stiffness hydrogels, they have higher expression of osteolytic factors and might possibly lead to higher metastatic load. This aligns with a study by Watson et al. that reports progressive osteolysis and increased osteolytic lesions when cells pre-conditioned in higher stiffness were injected (intra ventricular injection) in mouse models compared to cells pre-conditioned in low stiffness ^24^.

**Figure 7.**
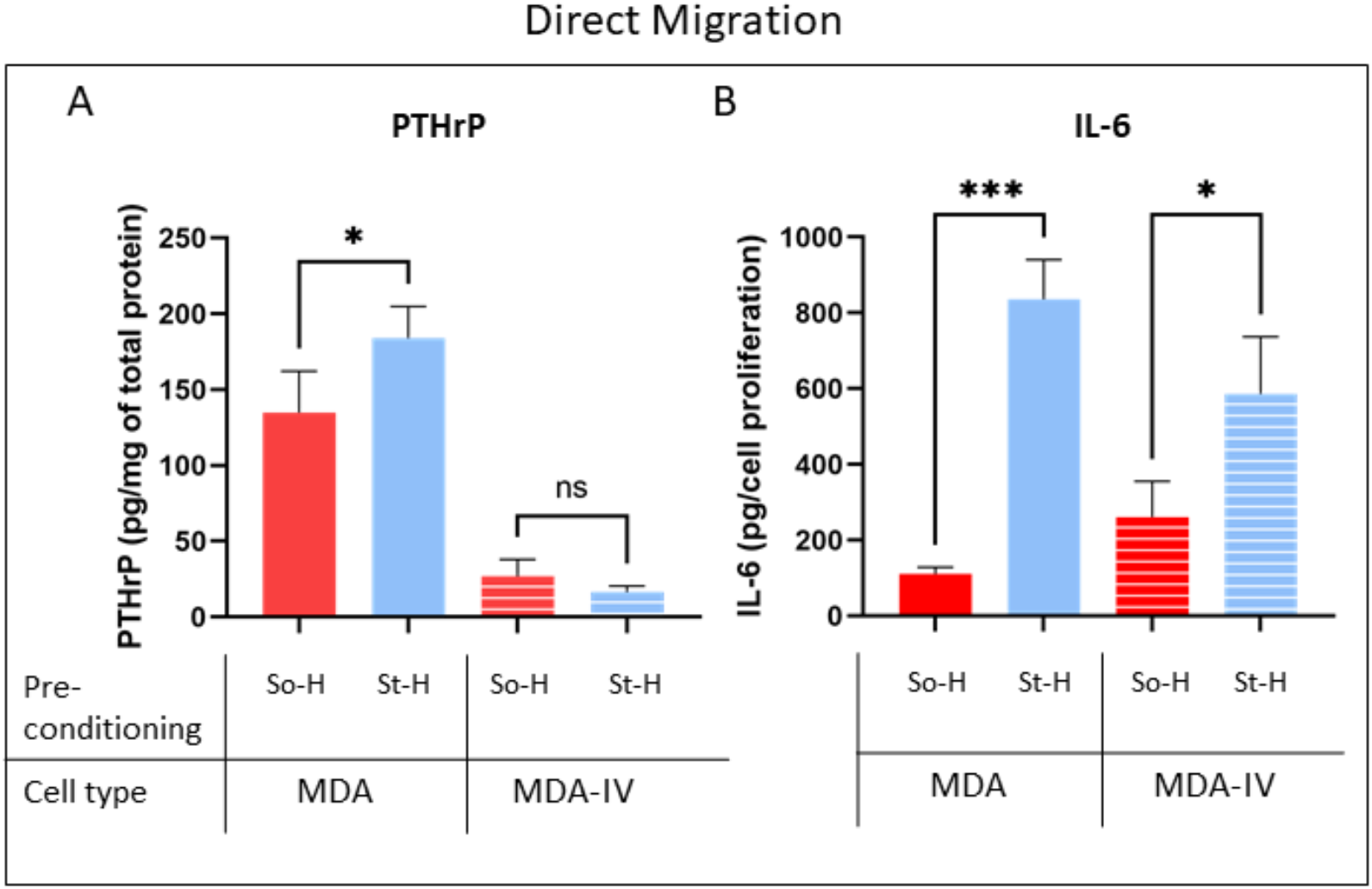
Direct migration model. **(A)** Bar graphs representing PTHrP expression and **(B)** IL-6 release in MDA-MB 231 (referred to as MDA in the graph) and MDA-IV cells. Results are normalized against the number of cells that have migrated to the biohybrid PCL HA scaffolds from So-H or St-H hydrogels. Data is represented as mean and SD of n=2 scaffolds and N=3 independent experiments. P-values represented as *p ≤ 0.05, **p ≤ 0.01, ***p ≤ 0.001, ****p ≤ 0.0001).

Overall, this study and the results presented here suggest that high primary tumor stiffness is linked with increased osteolysis. Specifically, our results suggest that expression of osteolytic factors could be a better indicator of bone metastatic load than cell migration /motility.

### 3.5 Response of MDA-IV cells in *Direct* and *Indirect migration model*

MDA-IV cells, a bone homing variant of MDA-MB 231, are known to exclusively metastasize to bone in vivo but show no difference in their colonization efficiency with respect to parental/original MDA-MB 231 ^44^. In this study, intrinsic PTHrP expression was found to be overall reduced in MDA-IV cells compared to parental MDA-MB 231 cell line when cultured in 2D (Figure 6A). On the contrary, when MDA-IV cells are cultured in *biohybrid PCL scaffolds*, PTHrP expression increased (both -/+ TGF-β supplementation), resembling what was observed with MDA-MB 231 (Figure 6A) and confirming that bone ECM induces expression of osteolytic factors in this cell line as well. Surprisingly, in *direct migration model*, expression of PTHrP in MDA-IV was low and did not vary as function of stiffness (Figure 7A). These results could suggest that while PTHrP is an important factor in bone colonization, it might not be as important to localize from breast to the bone. In fact, studies on patient tumors indicate that positive PTHrP expression in primary tumor is linked to lower bone metastasis and better patient survival ^45,46^. This could explain why MDA-IV cells have intrinsically low PTHrP expression than parental MDA-MB 231 when tested in vitro.

No difference was found in IL-6 release between MDA-IV and MDA-MB-231 cells when cultured in 2D TCP. *indirect migration models* showcased that levels of IL-6 increased in MDA-IV when cultured in PCL scaffolds, and were similar to the response of MDA-MB 231 pre-conditioned in the stiffer microenvironment (St-H, Figure 6B). This pattern was also observed in MDA-IV cultured in the *direct migration models*, where IL-6 release was found to have 2-fold increase in cells cultured in stiff hydrogels (Figure 7B). These results suggest that IL-6 might be an important marker in both localization and in remodeling of bone ECM during metastasis formation. Also, MDA-MB 231 evidenced a link with increasing hydrogels stiffness and high fold change of IL-6 (8 times) than PTHrP (1.4 times) in *direct migration model*. Importance of IL-6 cytokine in initiation of breast cancer invasion and metastasis ^47,48^ as well as its involvement in bone metastasis and osteolytic activity has been documented ^21,49^, which aligns with our findings for IL-6.

Results obtained in *direct* and *indirect migration models* suggest that it is possible to de-couple two different properties of breast-to-bone metastasis namely, localization and osteolytic/remodeling potential, whilst also examining the importance of different markers in these processes.

## Conclusions

Interactions between ECM and cancer cells direct phenotypes and matrix remodeling during tumor progression. In this study, engineering such matrix changes in vitro was found to be useful in isolating specific pattern of ECM properties and understand their impact on biological processes. Here, we used specific 3D in vitro models to elucidate effects of primary tumor matrix stiffness and gelatin-related adhesion motifs on various dimensions of breast cancer metastasis (i.e. adhesion, migration, 3D invasion, 3D metastatic site response). Stiffer primary tumor microenvironments (compressive moduli > 6 kPa) were found to be essential in inducing migratory and invasive phenotypes in MDA-MB 231 cells. In this study we focused on recreating aspects of breast-to-bone metastasis in vitro, and combined a 3D model matching human breast cancer tissue properties with a 3D model matching the properties of bone ECM. Two models were set-up to study metastatic onsets from breast-to-bone and focused on bone localization vs bone remodeling. The *indirect migration model* evidenced that both primary tumor stiffness and secondary site ECM composition are essential in regulating expression of osteolytic factors PTHrP and IL-6, and hence could affect osteolytic bone remodeling. We found that increased osteolytic potential in bone metastatic site is linked to stiffer primary tumor microenvironments. The *direct migration model* evidenced that high stiffness in the breast tumour is linked to high IL-6 activity, an important factor for metastasis initiation as well as osteolysis.

Engineered in vitro models, in accordance with previous in vivo study, were able to confirm that high primary tumor stiffness is linked to increased migration, invasion and osteolytic factors expression in breast cancer cells. Moreover, the models allowed to decouple different ECM properties and elucidate effect of both primary tumor ECM and secondary site ECM in breast-to-bone metastasis. Further inclusion of primary/patient-derived cells of known metastatic status in the proposed engineered in vitro models would be useful to predict possibility of bone metastasis and used as a platform to test therapeutic efficacy / tailor personalized treatments.

