## Supplementary material for "Invasion and secondary site colonization as a function of in vitro primary tumor matrix stiffness": Figure S1

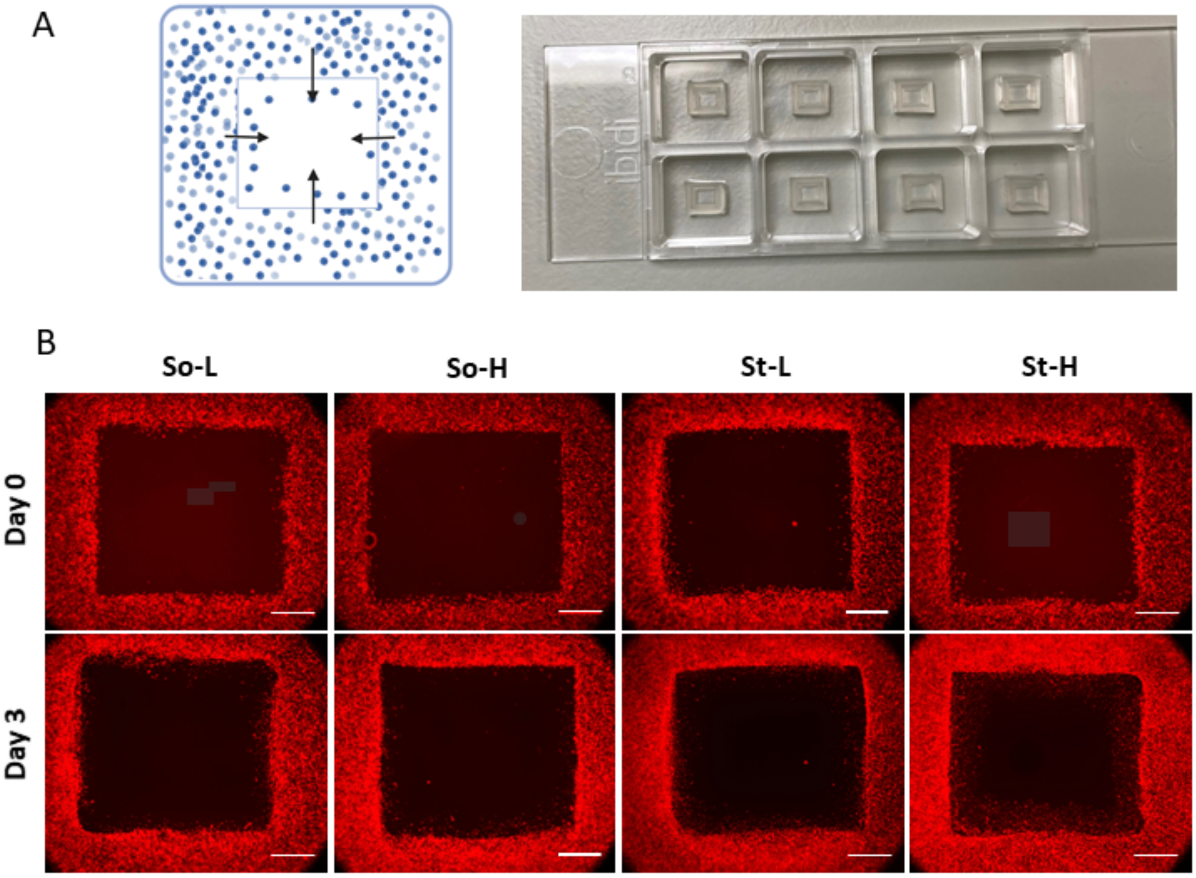


**Figure S1. 3D Collagen invasion assay. (A)** Schematic of invasion assay and representative image of inserts in 8-well imaging slides. **(B)** Cells stained with Cytopainter red were imaged at day 0 and day 3 of the assay (Scale bar- 1000 µm). Cells conditioned within four hydrogel groups of So-L, So-H, St-L and St-H were used in this assay. Insets and brightened image of day 3 invasion in acellular region showed in figure 4.
