## Supplementary material for "Invasion and secondary site colonization as a function of in vitro primary tumor matrix stiffness": Figure S2

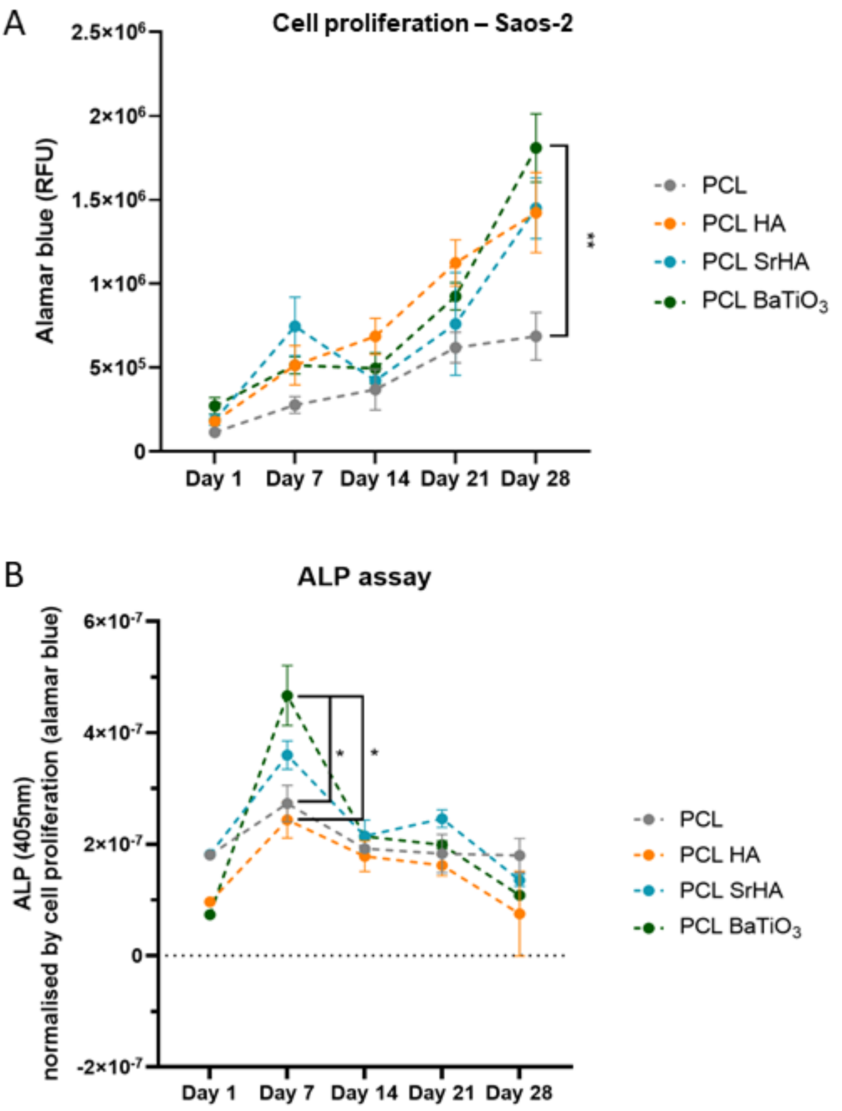


**Figure S2. (A)** Saos-2 proliferation as measured by Alamar blue assay and **(B)** ALP activity measured in Saos-2 cells when cultured in composite PCL scaffolds- PCL, PCL HA, PCL SrHA and PCL $BaTiO_3$. Data is plotted as a line graph from day 1 to day 28 with a 7-day interval (average and SD of three independent experiments. P-values represented as $*p \leq 0.05$, $**p \leq 0.01$, $***p \leq 0.001$, $****p \leq 0.0001$
