## Supplementary material for "Invasion and secondary site colonization as a function of in vitro primary tumor matrix stiffness": Figure S3

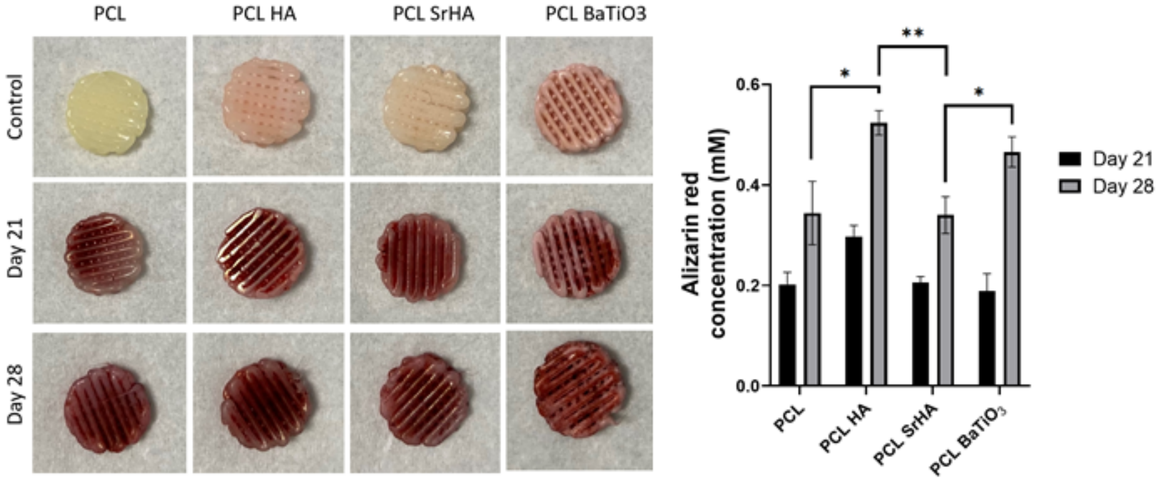


**Figure S3. Calcium deposition analysis.** PCL composite scaffolds stained with Alizarin stain on day 21 and day 28 of culture with Saos-2 cells (left). Control consisted of staining scaffolds with no cells cultured. Quantification of Alizarin stain on day 21 and day 28 and data represented as average and SD of three independent experiments (right). P-values represented as *$p \leq 0.05$, **$p \leq 0.01$, ***$p \leq 0.001$, ****$p \leq 0.0001$
