## Supplementary material for "Invasion and secondary site colonization as a function of in vitro primary tumor matrix stiffness": Figure S4

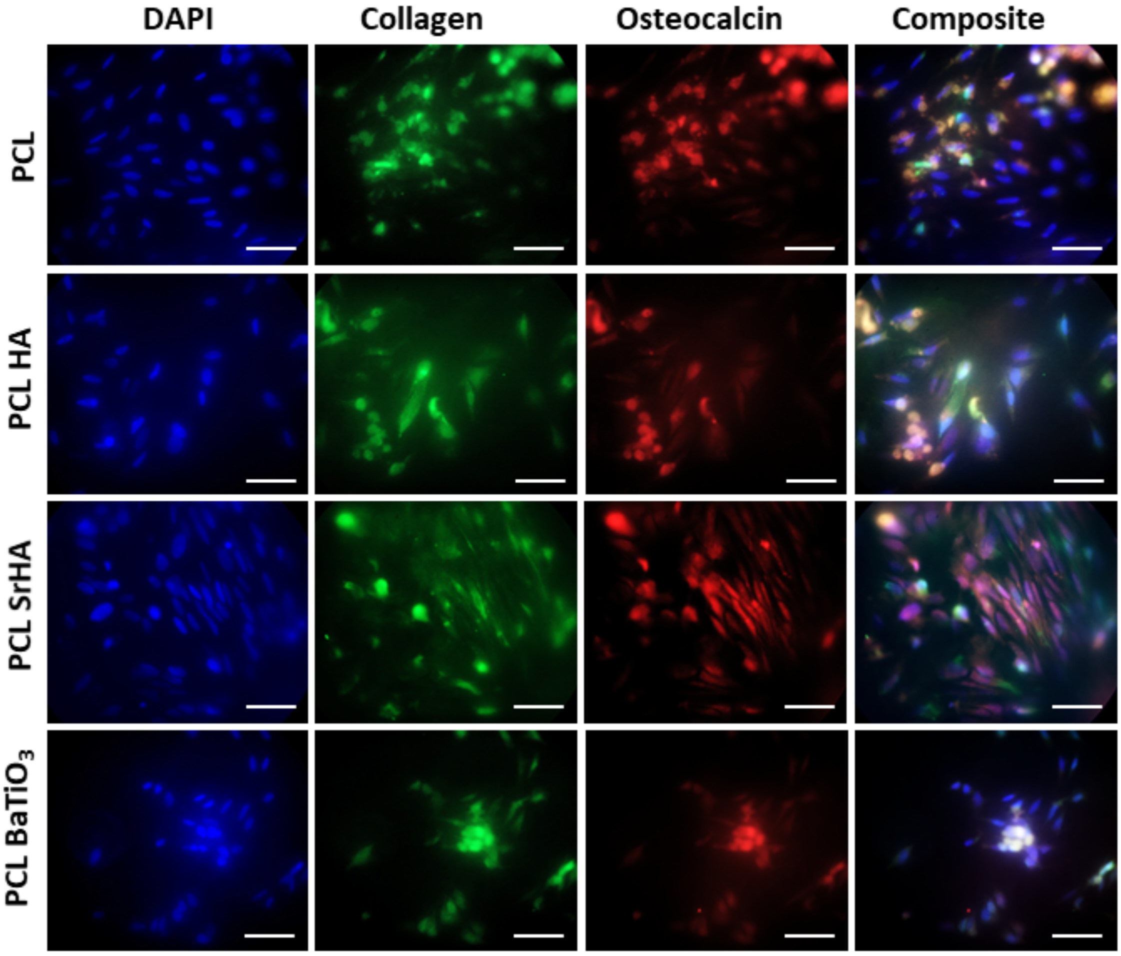


**Figure S4 Collagen and osteocalcin deposition.** Saos-2 cells were stained with DAPI, Collagen Ab and osteocalcin Ab on day 28 of culture in composite PCL scaffolds (Scale bar- 50 µm). The presence of punctate staining near nucleus of cells is considered to be ECM deposited by cells.
