## Supplementary material for "Invasion and secondary site colonization as a function of in vitro primary tumor matrix stiffness": Figure S5

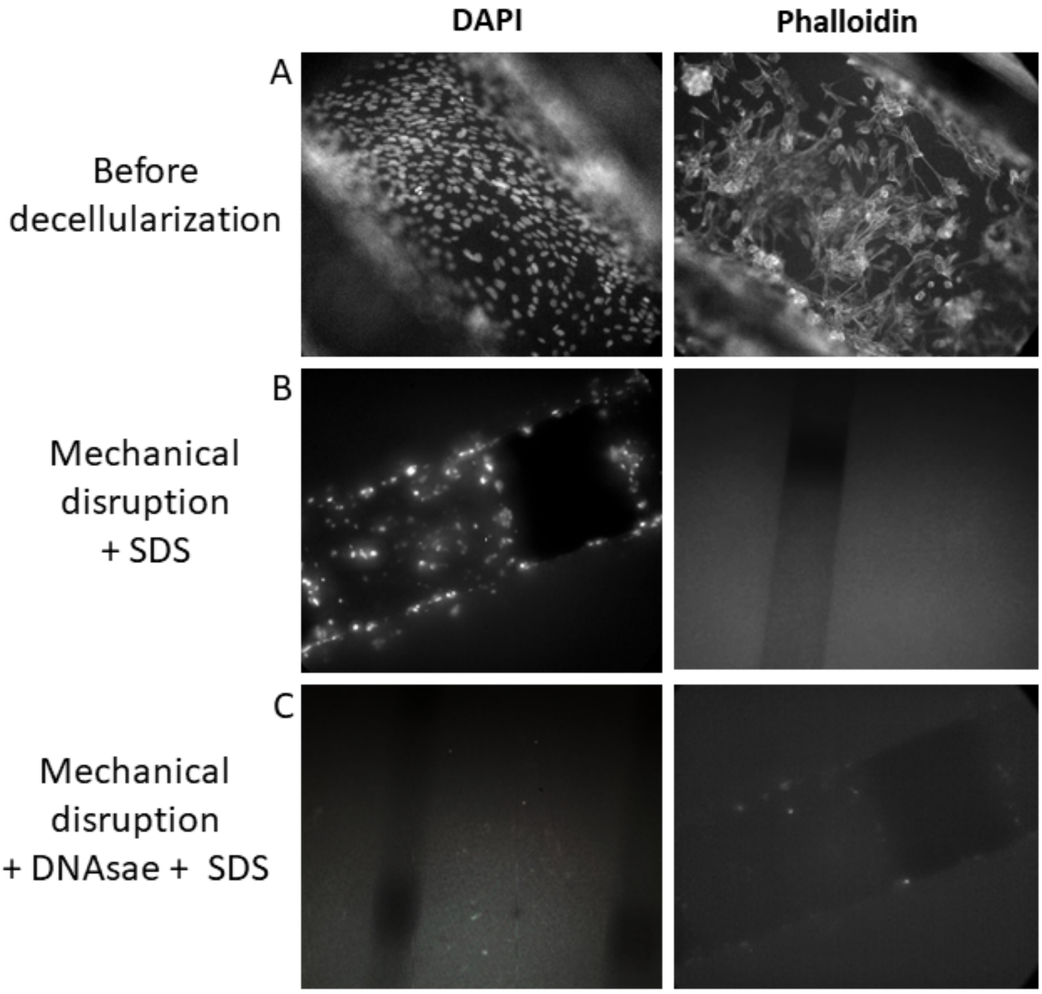


**Figure S5. Standardisation of decellularization process in composite PCL scaffolds.** DAPI and Phalloidin stained PCL scaffolds **(A)** before decellularization, (B) after mechanical lysis of cells and 0.05% SDS wash, and **(C)** after mechanical lysis with SDS and 1mg/mL DNAase.
