## Supplementary material for "Invasion and secondary site colonization as a function of in vitro primary tumor matrix stiffness": Figure S6

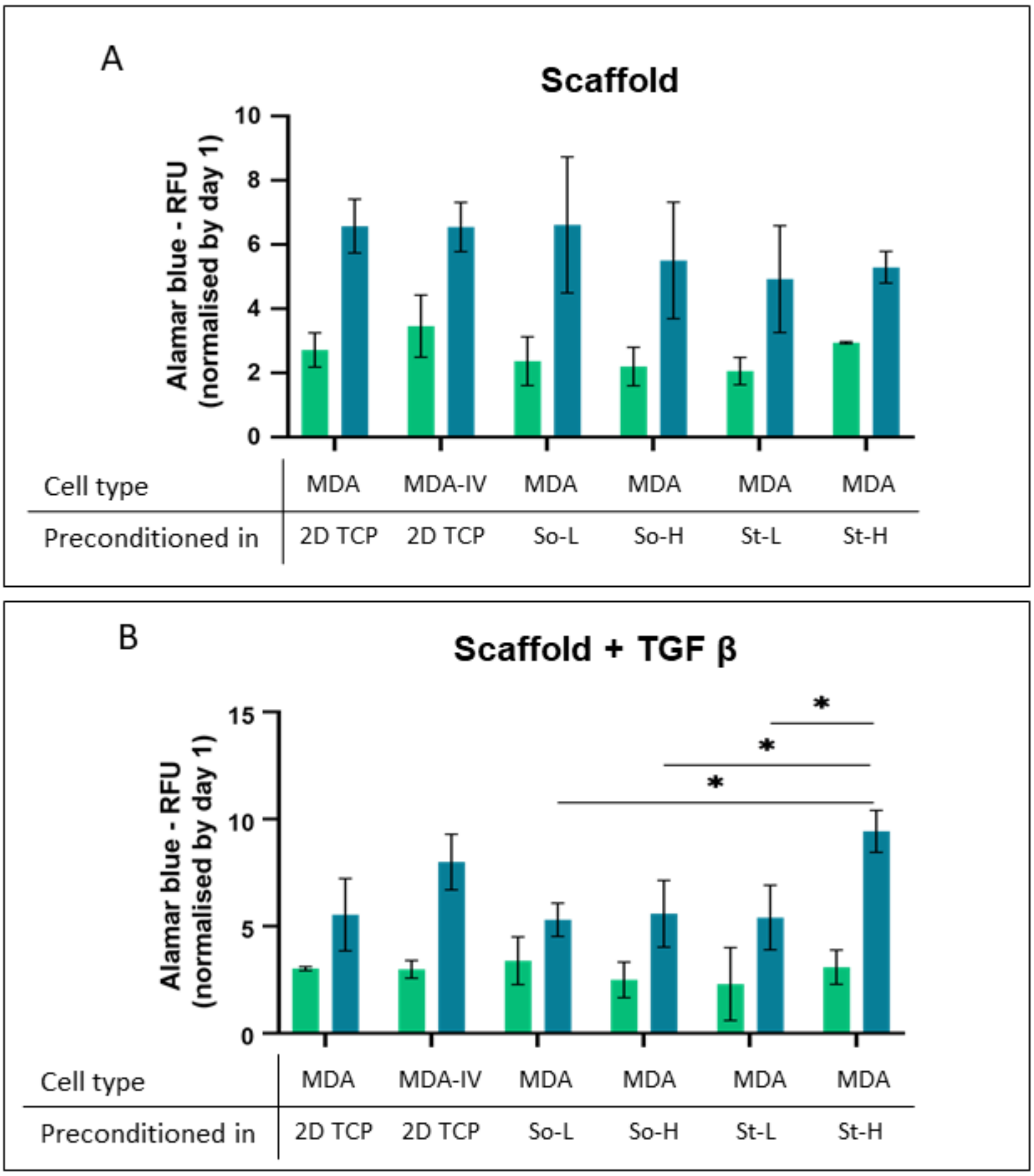


**Figure S6.** ***Indirect migration model*** **- Cell proliferation of breast cancer cells in biohybrid PCL scaffolds.** Alamar blue assay readings on day day 3 (green) and day 7 (blue) of MDA-MB 231 (represented as 'MDA') and MDA-IV cells pre-conditioned in either 2D TCP plates or alginate-gelatin hydrogels and plated on **(A)** Scaffolds and **(B)** Scaffolds + 5ng/mL TGF-β. Data is represented as average and SD of three independent experiments (right). P-values represented as $*p \leq 0.05$.
