## Supplementary material for "Invasion and secondary site colonization as a function of in vitro primary tumor matrix stiffness": Figure S7

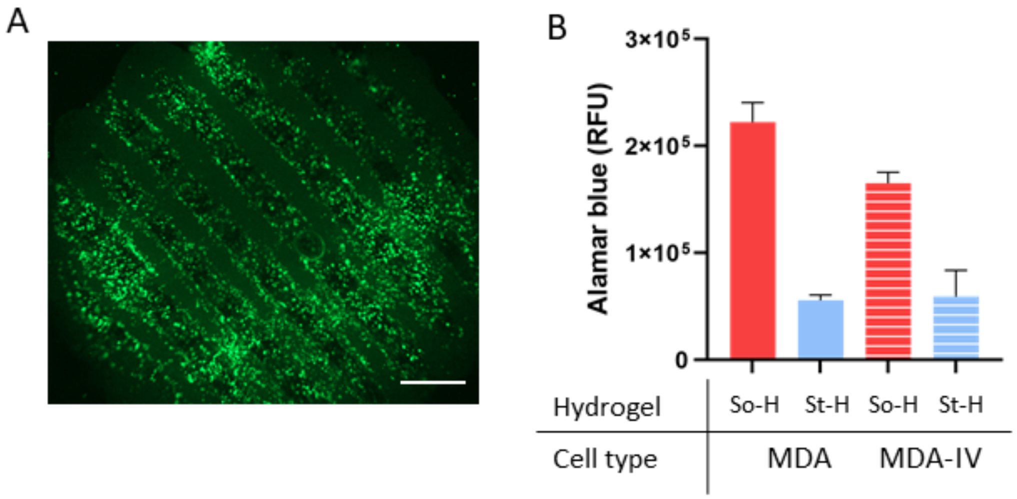


**Figure S7. Migration quantified in *direct migration model*. (A)** Cells migrated from alginate hydrogels to biohybrid scaffolds stained with green Live stain (ethylene homodimer) at day 7 (Scale bar- 1000 µm). **(B)** Quantification of migrated cells (MDA or MDA-IV) from So-H or St-H hydrogels to biohybrid PCL scaffold with Alamar blue assay at Day 7.
